# Bioinformatic identification of ClpI, a class of Clp unfoldase in Actinomycetota

**DOI:** 10.1101/2023.02.02.526855

**Authors:** Jialiu Jiang, Karl R. Schmitz

## Abstract

All clades of bacteria possess Hsp100/Clp family unfoldase enzymes that contribute to aspects of protein quality control. In Actinomycetota, these include ClpB, which functions as an independent chaperone and disaggregase, and ClpC, which cooperates with the ClpP1P2 peptidase to carry out regulated proteolysis of client proteins. We initially sought to algorithmically catalog Clp unfoldase orthologs from Actinomycetota into ClpB and ClpC categories. In the process, we uncovered a phylogenetically distinct third group of double-ringed Clp enzymes, which we term ClpI. ClpI enzymes are architecturally similar to ClpB and ClpC, with intact ATPase modules and motifs associated with substrate unfolding and translation. While ClpI possess an M-domain similar in length to that of ClpC, their N-terminal domain is more variable than the strongly conserved N-terminal domain of ClpC. Surprisingly, we identified separate sets of ClpI sequences that possess or lack the LGF-motifs required for stable assembly with ClpP1P2. In species where they occur, we suggest that ClpI enzymes provides additional pathways and points of regulatory control over protein quality control programs, supplementing the conserved roles of ClpB and ClpC.

## INTRODUCTION

All bacteria possess double-ringed Hsp100/Clp family AAA+ (ATPases Associated with various cellular Activities) enzymes that harness chemical energy from ATP to unfold client proteins. In Actinomycetota (synonym Actinobacteria), these include the enzymes ClpB and ClpC, which participate in distinct aspects of protein quality control. ClpB operates as a chaperone and disaggregase, remodeling and releasing client proteins to refold in the cytosol (1–3). ClpC functions as the unfoldase component of the ClpCP1P2 protease (4–6). Target proteins unfolded by ClpC are translocated into the degradation chamber of the associated ClpP1P2 peptidase for degradation (7, 8). Although the specific roles of ClpCP1P2 in Actinomycetota are poorly defined, this protease has emerged as a promising antibacterial target in *Mycobacterium tuberculosis*, and multiple compounds are known to kill *M. tuberculosis* by disrupting ClpCP1 P2 activity (9–16).

ClpB and ClpC share a common architecture and operating principles. Both enzymes consist of a family-specific N-domain followed by two AAA+ ATPase modules, and both function as homomeric hexamers with a central axial channel (17–20). Protein substrates are engaged by axial loops within the channel (21, 22). ATP hydrolysis events in individual ATP modules drive conformational changes in the ring that apply an unfolding force to gripped substrates (21, 23). The interaction between ClpC and the ClpP1P2 peptidase is stabilized by flexible LGF-loops present on the underside of the unfoldase, which dock into hydrophobic pockets on the surface of the peptidase (**Fig.1A**) (7, 24). ClpB lacks LGF-loops and thus cannot collaborate with ClpP1P2 to carry out ATP-dependent proteolysis (3, 25). Indeed, the presence or absence of LGF-loops serves as a useful sequence marker for discriminating ClpB and ClpC sequences.

**Figure 1.**
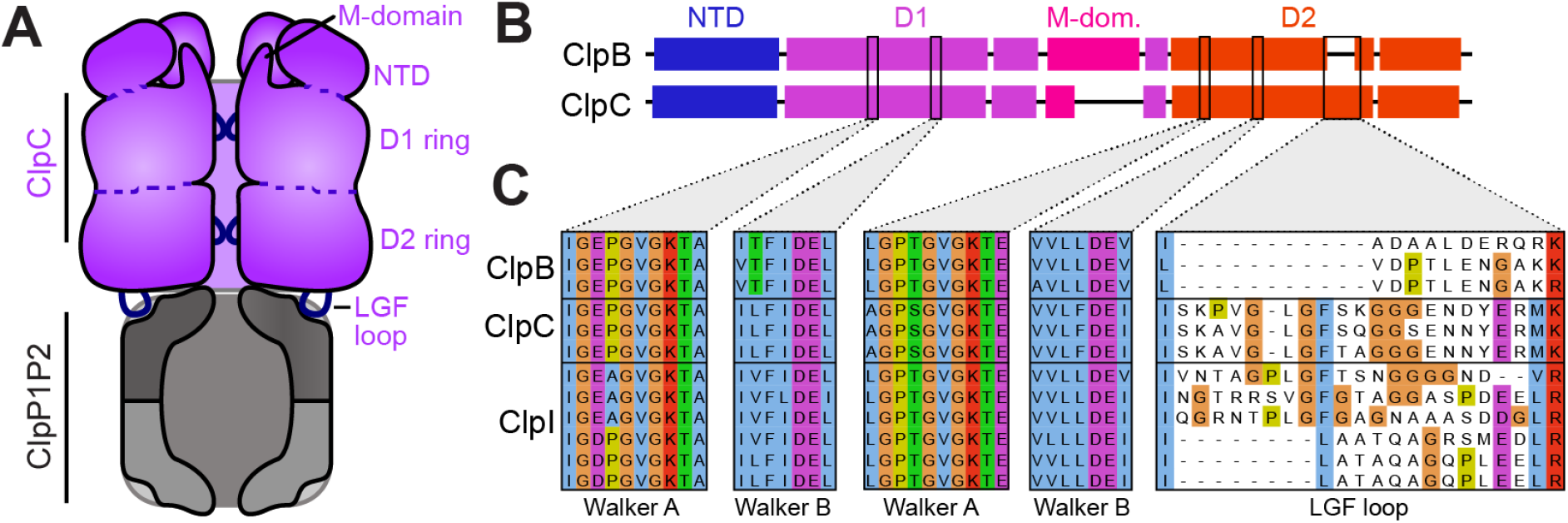
Clp enzyme architecture and domain organization. **A**) A cartoon of ClpC illustrates notable features of Clp enzymes. In the context of proteolysis by ClpCP1P2, client substrates are recognized by ClpC, unfolded, and translocated through the axial pore into the peptidase degradation chamber. **B**) [Label Walter motifs, and LGF loop] ClpB and ClpC proteins contain a family-specific N-terminal domain (NTD), followed by two AAA+ ATPase modules (D1 and D2) with conserved Walker A and B motifs (WA and WB). The D1 ATPase module is interrupted by a helical M-domain that projects upward towards the NTD. ClpC enzymes additionally possess an LGF loop that makes stabilizing contacts with the ClpP2 ring of the peptidase. **C**) Sequence alignment of select ClpC orthologs from Actinomycetota shows strong conservation of Walker A and B motifs. Notably, a subset of sequences possesses a short M-domain and distinct LGF loop regions, with or without identifiable LGF-like motifs.

We initially sought to discriminate Actinomycetota ClpB and ClpC enzymes based on characteristic sequence features, as a prerequisite for analyzing the unique patterns of sequence conservation among ClpC orthologs. In the process, we uncovered a group of ClpC/B paralogs with intermediate characteristics, which we term ClpI enzymes (**Fig. 1C**). Bioinformatic analyses reveals that ClpI sequences possess conserved features associated with ATP hydrolysis and unfoldase activity, but are evolutionarily distinct from both ClpC and ClpB. Notably, ClpI sequences occur with and without LGF-loops, suggesting that individual lineages of ClpI enzymes have evolved to work either as independent unfoldases or to proteolyze client proteins in cooperation with ClpP1P2. Our findings expand on the known diversity of AAA+ unfoldases, and reveal new points of regulation by which species within Actinomycetota can modulate protein quality control.

## METHODS

### Sequence analysis of Actinomycetota ClpC paralogs

Ortholog sequences were collected using NCBI BLAST (26) and the HMMER search algorithm (27). The BLAST bit-score was used to assess the similarity of hits to reference sequences. 2D scatterplots illustrating the similarity of dataset members to reference sequences were constructed using the Python 3 matplotlib package (28, 29). Each data point in the resulting plots corresponds to the sequence of a single ClpC paralog.

As an independent assessment of similarity, principal component analysis datasets was used as a dimension reducing method (30). Alignment scores of each sequence were determined and expressed as principal components with components equal to 10.

### Multiple sequence alignments and phylogenetic analysis

Multiple sequence alignments of ClpC, ClpB, or ClpI orthologs were created using ClustalW (31). To compare the ClpC/B/I sequence occurrence among phylogenetic groups, the taxonomic order associated with each hit was extracted from the Uniprot database using Biopython (32, 33). For phylogenetic analysis, 5 sequences were randomly selected from each taxonomic order (for orders with fewer than 5 representatives all sequences were included), sequences were aligned using CLUSTALW, and the resulting alignment was used to guide construction of a phylogenetic tree using ETE TOOLKIT (34, 35). Clusters with bootstrap values greater than 50% were defined as confirmed subgroups (36).

### Cross-alignment sequence comparison

Pre-calculated ClustalW alignments of individual ClpC/B/I paralogs were aligned to one another using the profile-profile method in MUSCLE (37). To illustrate positional conservation between top and bottom alignments, a 20-member profile array *P* of amino acid frequency *f* was constructed for each position in each alignment: *P* = (*f*_1_, …, *f*_20_) (38). For each residue *x* in an alignment, a BLOSUM62-derived similarity score was calculated by summing BLOSUM62 substitution scores for that residue (39), weighted by the profile frequency of substituted residues *y*.

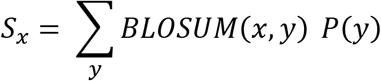

Alignments were represented as color coded bitmaps, with each position colored according to its score. A cross-alignment similarity score *S* was calculated for each position by iterating over all residue combinations (*x,y*) and summing BLOSUM62 scores weighted by the profile frequency in both the top (*P_t_*) and bottom (*P_b_*) frequency profile arrays.

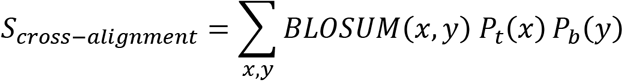

Positional cross-comparison scores were represented as a one-dimensional heatmap between alignments.

## RESULTS

### Actinomycetota possess a group of unusual Clp unfoldases

As an outgrowth of our interest in mycobacterial ClpC, we sought to assess the pattern of amino acid conservation across ClpC orthologs within Actinomycetota. We used the search algorithm HMMER (27) to collect Actinomycetota homologs of *Mycolicibacterium smegmatis* ClpC1 and aligned these using ClustalW (31). ClpC bears substantial sequence and structural homology to ClpB, and our search results consequently included numerous hits annotated as ClpB or ambiguously as Clp enzymes. We attempted to use known sequence features to definitively sort homologs into separate ClpC and ClpB categories. Both groups of enzymes contain conserved motifs associated with ATP hydrolysis and unfolding/translocation (**Fig. 1C**), but we note two major sequence features distinguish them: *i*) the M-domain that projects upward from the D1 ATPase module is shorter in ClpC1 (~ 20 aa) than in ClpB (~ 90 aa), and *ii*) ClpC, but not ClpB, contains an LGF-loop with a Leu-Gly-Phe or similar motif that stabilizes binding to the surface of the ClpP1P2 peptidase (8, 24, 40). Based on these features, we were able to categorize the majority of sequences as either ClpC or ClpB orthologs. However, a subset of sequences possessed short ClpC-like M-domains and either lacked LGF-loops or lacked identifiable LGF-like motifs. These unusual sequences had higher homology to ClpC and ClpB than to other AAA+ enzymes. From these sequence features alone, it was unclear whether intermediate sequences were atypical ClpC enzymes, atypical ClpB enzymes, or an entirely separate category.

To understand the relationship between atypical sequences and ClpC/ClpB groups, we used NCBI BLAST to assess the similarity of each entry in our Actinomycetota dataset to *M. smegmatis* ClpC1 and ClpB reference sequences. BLAST bit-scores to each reference were plotted as a two-dimensional scatter plot (**Fig. 2A**). The majority of sequences lie in two off-diagonal clusters: one with higher similarity to ClpC (**Fig. 2A** yellow box; 1294 sequences) and one with higher similarity to ClpB (**Fig. 2A** pink box; 1162 sequences). Inspection of individual hits in these clusters confirmed that they comprise canonical ClpC and ClpB sequences, based on sequence features and annotations. However, a subset of sequences occupied a distinct third cluster, with moderate but approximately equal similarity to ClpC and ClpB references (**Fig. 2A** blue box; 451 sequences). Importantly, sequences clustered into this third group based on overall sequence similarity to ClpB and ClpC references, rather than on specific sequence features. The intermediate cluster included the outlier sequences noted above that lack LGF-motifs, but also included a number of sequences with identifiable LGF-motifs (**Fig.1C**). Given their intermediate similarity to ClpC and ClpB references, we termed the proteins in this third group ClpI enzymes. ClpI subtypes with LGF-motifs were termed ClpIa, and those without LGF-motifs were labeled ClpIb.

**Figure 2.**
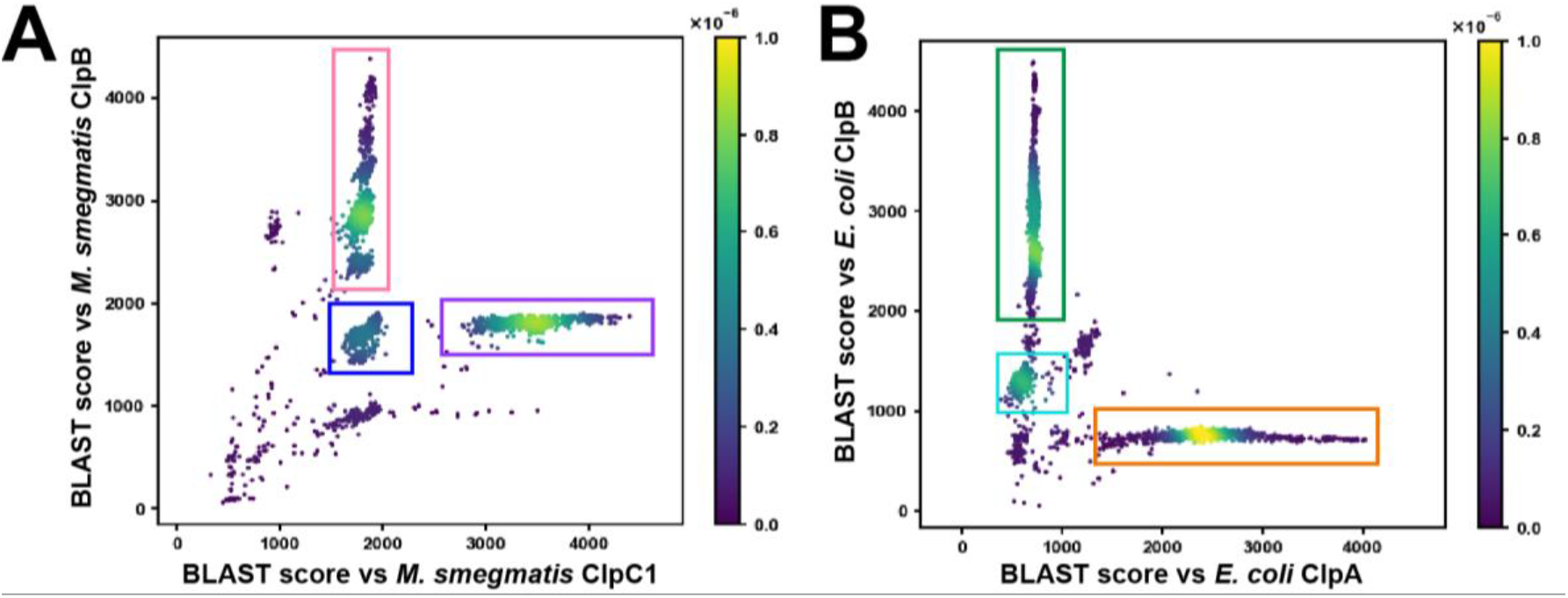
BLAST analysis of ClpC/B/A orthologs. **A)** Actinomycetota ClpC orthologs were compared to *Mycolicibacterium smegmatis* ClpB (Uniprot ID: A0QQF0_MYCS2) and ClpC (CLPC1_MYCS2) reference sequences. The *x* axis represents the BLAST bit-score resulting from alignment of dataset sequences with the ClpC reference, while the *y* axis represents the BLAST bit-score associated with alignment to the ClpB reference. The density of hits in a region of the plot is shown as a heatmap. Typical ClpC orthologs cluster within the purple box, typical ClpB orthologs within the pink box, and atypical sequences with intermediate features cluster within the blue box. **B)** Proteobacterial ClpA orthologs were compared to *E. coli* ClpA (CLPA_ECOLI) and ClpB (CLPB_ECOLI) references as in A. Conventional ClpA sequences lie within the orange box, conventional ClpB sequences lie within the green box, and orthologs of ClpV, a translocase associated with Type IV secretion systems, occupy the cyan box.

For comparison, we performed an equivalent analysis of proteobacterial Clp enzymes. HMMER was used to compile proteobacterial homologs of *E. coli* ClpA, which were plotted based on BLAST similarity to *E. coli* ClpA and ClpB references (**Fig. 2B**). The majority of hits appeared in either ClpA (orange box) or ClpB (green box) groups, with only a handful of sequences lying on the diagonal in two sparse clusters. A minor cluster located below the main ClpB group contained sequences of the Type VI secretion system ATPase ClpV/TssH (cyan box), which possess an M-domain but generally lack LGF-loops (41). Most other on-diagonal sequences were shorter than ClpA or ClpB, and thus possibly fragmentary genes or the result of mis-annotated translational start sites. The overall clustering pattern is in agreement with the expectation that ClpA and ClpB are the predominant groups of double-ringed Clp enzymes in proteobacteria (42). Moreover, the clustering of ClpV sequences validates this approach for identifying distinct ortholog subtypes, and makes it unlikely that the ClpI cluster is an artifact of the analysis method.

As an independent assessment of sequence clustering, we combined sets of sequences and subjected them to principle component analysis (PCA) (30). PCA plots consistently sorted ClpI sequences into groups separate from ClpB and ClpC (**Fig. S1**). ClpIa and ClpIb subtypes were more similar, and only partially resolved into separate clusters.

### ClpI orthologs occur in a subset of orders in Actinomycetota

ClpC, ClpB and ClpI sequences were not equally abundant within the set of ClpC homologs identified by HMMER. ClpC and ClpB made up the majority of the dataset with similar individual abundance (~40% and ~44% of total, respectively), while ClpI sequences comprised only ~16% of total. The lower prevalence of ClpI suggests that these are present only in a subset of clades.

To establish the distribution of ClpI enzymes across phylogenetic lineages, we binned ClpB, ClpC, and ClpI sequences from our dataset by taxonomic order (**Table 1**). ClpB and ClpC sequences were found in all 20 observed orders, suggesting that both enzymes play a critical role in cellular function and are evolutionary conserved across taxa. Interestingly, many individual orders contained unequal numbers of ClpB and ClpC sequences. For example, in Streptosporangiales we found 75 ClpB orthologs but only 50 ClpC orthologs. Conversely, Micrococcales contained 305 ClpB and 334 ClpC orthologs. As expected from the lower overall abundance, ClpI sequences were present in only 13 orders. In orders that possess ClpI orthologs, the number of ClpI sequences was generally lower than either ClpB or ClpC, suggesting that ancestral ClpI enzymes were lost in some lineages, and thus play a less critical role in cellular physiology. The prevalence of ClpIa and ClpIb subtypes within orders also varied. In most cases, (eg, Pseudonocardiales, Micromonosporales, Micrococcales) ClpIa sequences outnumbered ClpIb. Only in Mycobacteriales and Streptosporangiales were the two subtypes similarly abundant, although still less prevalent than ClpB or ClpC. Several orders (eg, Actinomycetales) possessed only a single ClpI sequence, which may have arrived by horizontal gene transfer. Notably, ClpI enzymes do not necessarily replace ClpB or ClpC enzmyes in a given species. We note species in which all four enzymes are clearly present – for example, *Rhodococcus sp. ABRD24* (ClpB Uniprot ID: ERC79_08130; ClpC: A0A4P6UCA9_9NOCA; ClpIa: A0A4P6UA74_9NOCA; ClpIb: A0A4P6UBT3_9NOCA).

**Table.1.** Occurrence of ClpB, ClpC and ClpI across Actinomycetota orders.

| **Order** | **ClpB** | **ClpC** | **ClpIa** | **ClpIb** |
| --- | --- | --- | --- | --- |
| **Pseudonocardiales** | 83 | 73 | 25 | 7 |
| **Actinomycetales** | 36 | 39 | 0 | 1 |
| **Kineosporiales** | 7 | 6 | 0 | 0 |
| **Nakamurellales** | 4 | 4 | 0 | 0 |
| **Jiangellales** | 6 | 6 | 1 | 0 |
| **Micromonosporales** | 64 | 58 | 50 | 5 |
| **Bifidobacteriales** | 46 | 36 | 0 | 1 |
| **Cryptosporangiales** | 1 | 1 | 0 | 0 |
| **Actinopolysporales** | 2 | 2 | 2 | 0 |
| **Frankiales** | 23 | 16 | 1 | 0 |
| **Catenulisporales** | 2 | 1 | 0 | 1 |
| **Propionibacteriales** | 113 | 96 | 0 | 0 |
| **Mycobacteriales** | 204 | 185 | 15 | 14 |
| **Acidothermales** | 1 | 1 | 0 | 0 |
| **Streptosporangiales** | 75 | 50 | 18 | 14 |
| **Glycomycetales** | 3 | 3 | 0 | 0 |
| **Streptomycetales** | 296 | 230 | 117 | 78 |
| **Coriobacteriales** | 1 | 0 | 0 | 0 |
| **Geodermatophilales** | 22 | 21 | 9 | 1 |
| **Micrococcales** | 305 | 334 | 88 | 3 |
| **Total** | 1294 | 1162 | 326 | 125 |
| **% total** | 44.51 | 39.97 | 11.21 | 4.30 |

To assess the similarity of Clp enzyme sequences within taxa, we grouped sequences by taxonomic order and plotted BLAST bit-scores generated against *M. smegmatis* ClpB (**Fig. 3**). ClpB orthologs showed a wide range of scores between 2000 and 4500. The diversity of ClpB scores, particularly within Mycobacteriales, reflects the fact that some ClpB sequences are highly similar to the reference, while more distantly related ClpB orthologs have more divergent sequence composition. However, there also appears to be consistent diversity in ClpB sequence composition across orders, which manifests as separate clusters spaced along the y-axis at scores of ~2300 and ~2600. Interestingly, all ClpC group members produced similar scores of ~2000, suggesting less variation of ClpC sequences across clades. ClpI sequences produced scores similar to ClpC, although both ClpIa and ClpIb subgroups exhibited greater variability in score than ClpC, suggesting lower sequence conservation.

**Figure 3.**
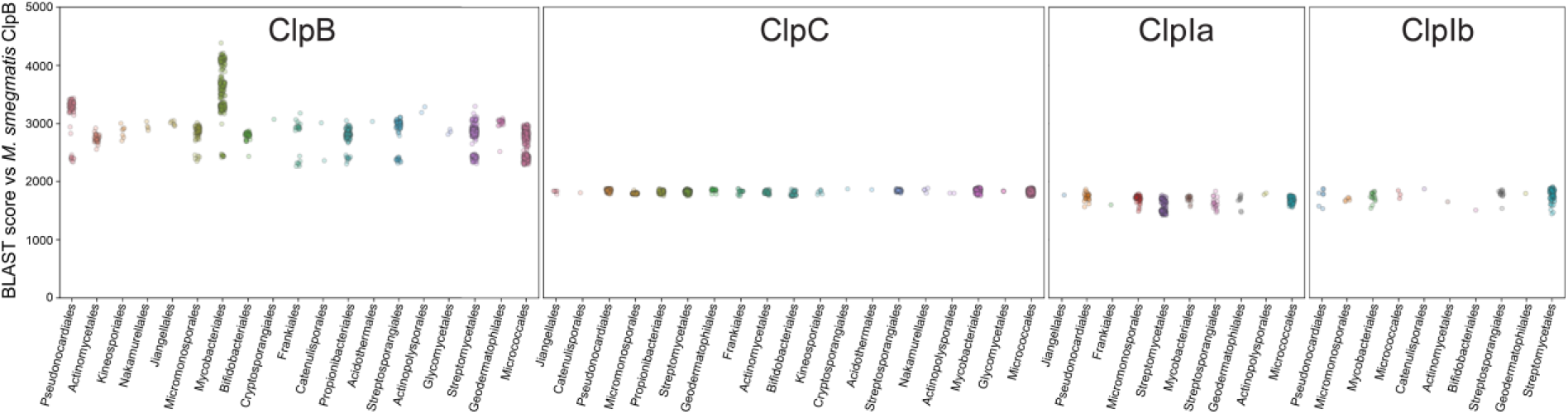
Sequence abundance and variability within orders. For each category of Clp unfoldases, sequences are grouped by taxonomic order and the BLAST bit-score calculated against *M. smegmatis* ClpB is plotted on the *y* axis.

### ClpIa and ClpIb subtypes are phylogenetically distinct

One curious characteristic of ClpI sequences is the presence of homologs with (ClpIa) and without (ClpIb) identifiable LGF-motifs. This heterogeneity stands in contrast to ClpC and ClpB groups, which uniformly possess or lack LGF-motifs, respectively. The existence of ClpI enzymes with and without LGF-motifs suggests either that ClpIa and ClpIb are evolutionarily distinct subgroups, or that LGF-motifs have been independently gained or lost over time in individual lineages.

To understand whether ClpIa and ClpIb subgroups represent separate evolutionary lineages, we performed phylogenetic analysis on a subset of ClpC, ClpB, and ClpI enzymes from each order within Actinomycetota (**Fig. 4**). ClpB sequences form a distinct phylogenetic group, as do most ClpC and ClpI enzymes, in agreement with NCBI BLAST analysis (**Fig. 2A**). (Bifidobacteriales ClpC and ClpI are an exception, clustering together in a branch between the major ClpC and ClpI divisions.) Interestingly, we observe that ClpIa and ClpIb subtypes sort into separate divisions within the main ClpI branch. This pattern suggests that duplication of an ancestral ClpI-encoding gene (or horizontal acquisition of an ancestral ClpI) occurred early in Actinomycetota evolution, allowing differentiation into ClpIa and ClpIb paralogs with distinct cellular roles.

**Figure 4.**
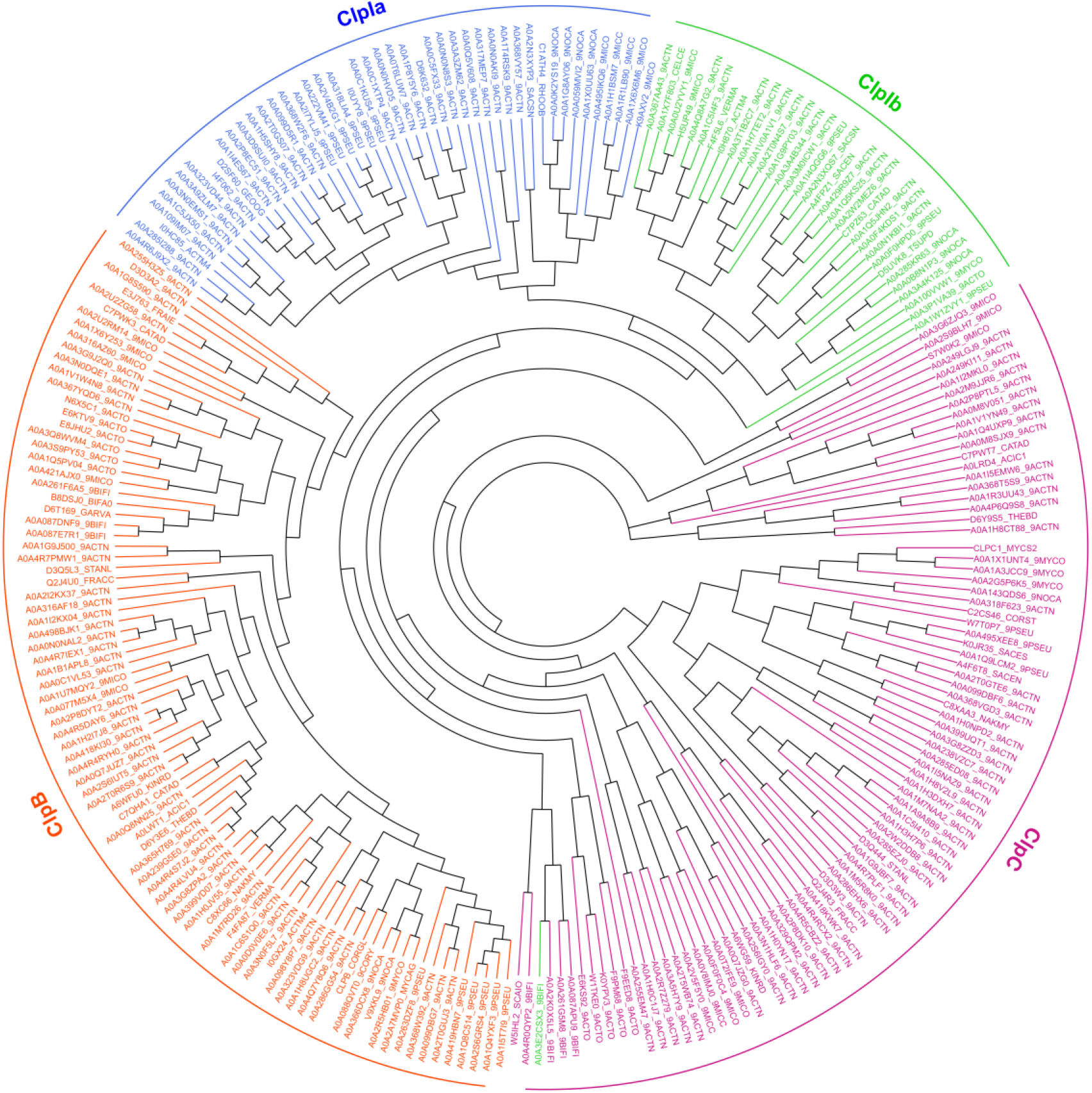
Phylogenetic analysis of ClpC paralogs. Distance methods were used to generate a phylogenetic tree of ~250 representative sequences of Actinomycetota ClpB (orange), ClpC (violet), ClpIa (blue), and ClpIb (green). Leaves are labeled with Uniprot accession IDs.

### Comparison of sequence conservation patterns among Clp unfoldases

To examine the sequence conservation within and across enzyme groups, we compared multiple sequence alignments generated for each group (**Fig. 5**). Comparing sequence conservation patterns across alignments, we found that the large and small subdomains of the D1 and D2 ATPase modules were similarly conserved across all enzyme groups. ClpB enzymes are distinguished by substantially longer M-domains than either ClpC or ClpI (**Fig. 5A,B**). The ClpB M-domain contains two sites that tolerate insertions of variable length, whereas the length of the ClpC M-domain is strictly conserved. The equivalent region in ClpI is similar in length to that of ClpC (**Fig. 5C**). Interestingly, M-domain insertions appear to be tolerated in ClpIb but not ClpIa enzymes (**Fig. 5D**), suggesting that a specific M-domain length is more important in the context of proteolysis than disaggregation.

**Figure 5.**
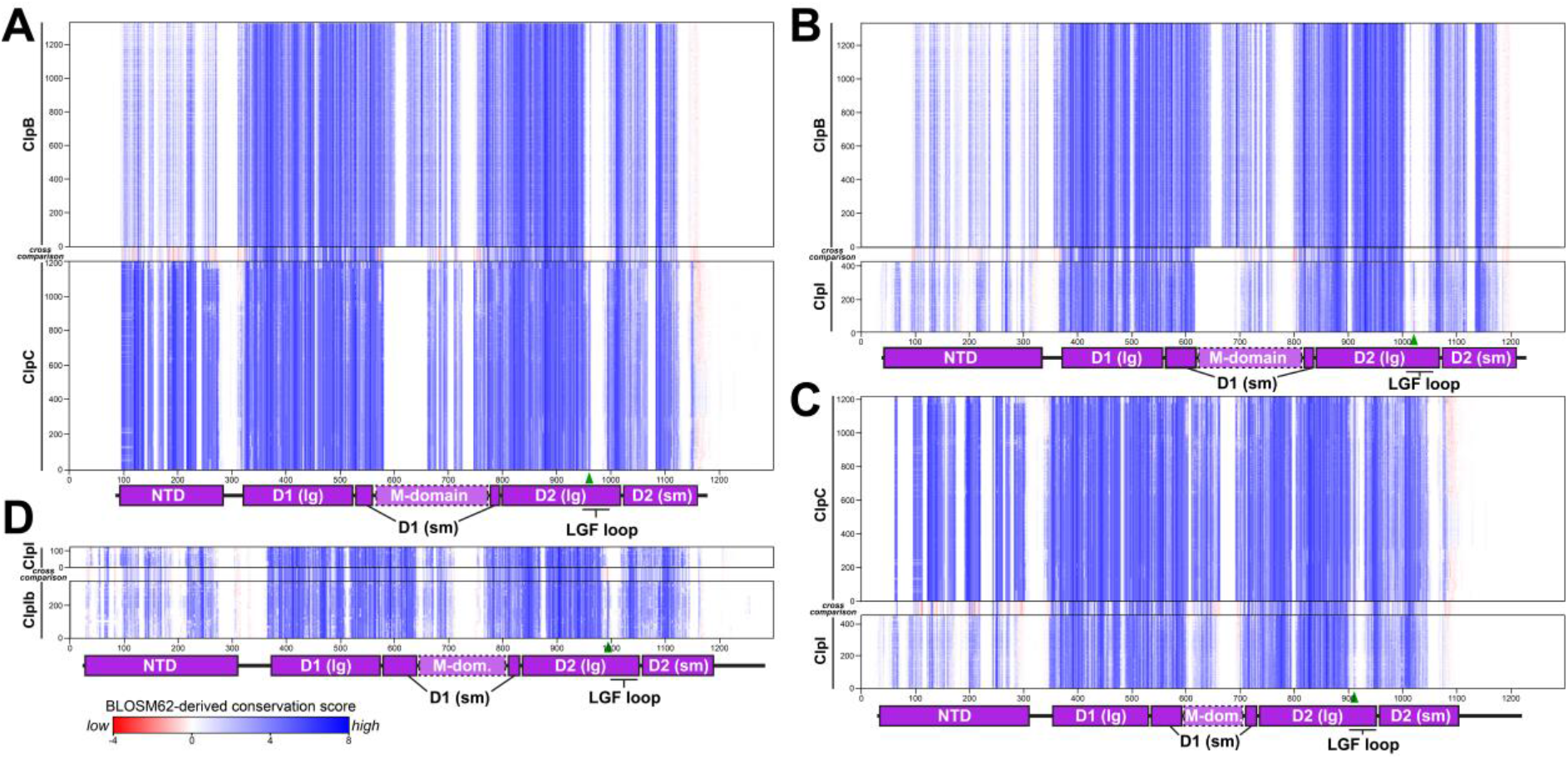
Comparison of sequence conservation patterns. Multiple sequence alignments of Clp unfoldase orthologs are shown as bitmaps with each residue represented as a single color-coded pixel. Residues are colored according to their similarity to the overall amino acid frequency at that alignment position. Pairs of sequence alignments are cross-aligned. A cross comparison heatmap illustrates the similarity in residue frequency at each position in the top and bottom alignments: **A**) ClpB is aligned to ClpI, **B**) ClpB to ClpI, **C**) ClpC to ClpI, and **D**) ClpIa to ClpIb. Strong conservation within or across alignments appears blue; low conservation appears red.

One stark difference among these enzyme groups is the extent of sequence conservation within the NTD. In both ClpB and ClpI, the NTD is less conserved than the ATPase modules (**Fig. 5B**), with strict conservation among only a subset of residues that form its folded core. (About 10% of ClpB sequences lack NTDs altogether, although these may be due to mis-annotated start sites.) Several sites in the ClpI NTD tolerate insertions, including the extreme N-terminus. By contrast, the ClpC NTD is a fixed length with high sequence conservation even outside of the folded core, comparable to the ATPase modules (**Fig. 5C**). This suggests that interactions made by the ClpC NTD, such as those with substrates, adaptors, or the M-domain, are both important for function and conserved across Actinomycetota. Conversely, sequence-specific functions of ClpB and ClpI NTDs are either less important or more diverse. Whereas M-domain length serves as a strong marker for ClpB sequences, the conserved NTD sequence is the strongest diagnostic feature distinguishing ClpC orthologs. Given the marked difference in NTDs between enzyme groups, we were curious whether the NTD alone drives bioinformatic partitioning of sequences into distinct ClpB, ClpC, and ClpI groups. To test this, we truncated NTDs from our dataset and performed BLAST analysis against *M. smegmatis* ClpB and ClpC1, as in **Figure 2A**. Truncated sequences similarly clustered into ClpB, ClpC and ClpI groups (**Fig. S2**), demonstrating that characteristics beyond the NTD support the existence of an independent ClpI cluster.

Interestingly, ClpC and ClpI NTDs are followed by a region of low conservation and variable length (**Fig. 5B,C**), consistent with a flexible linker leading to the large subunit of the D1 ring. Indeed, the NTD was not resolved in recent cryo-EM structures of *M. tuberculosis* ClpC1 (7), presumably due to its conformational flexibility with respect to the enzyme core. Conversely, the linkage between NTD and D1 is shorter in ClpB and lacks insertions (**Fig. 5A**), suggesting lower conformational flexibility between the ClpB NTD and the body of the unfoldase. Indeed, AlphaFold2 predictions of ClpB enzymes from Actinomycetota members (eg, model AF-P9WPD1-F1 of ClpB from *M. smegmatis* H37Rv) show most of this region as packed against the NTD.

We examined sequence alignments to identify positions that were strongly conserved within individual enzyme groups, but differed in residue identity between groups. These sites of contrasting conservation generally correlate with red lines in the cross-comparison strips shown in **Figure 5**. Positions with notable enzyme-specific amino acid identities are listed in **Table S1**, and may be useful in sorting uncharacterized Actinomycetota Clp enzymes into ClpB/C/I groups.

### Differences in NTD surface conservation

Our original motivation for classifying Clp unfoldases was to understand sequence conservation patterns among ClpC orthologs. Because the degree of overall NTD conservation varied among ClpB/C/I enzymes, we used Consurf (43–45) to map sequence conservation onto models of the NTD from each group and compared the results (**Fig. 6**). In agreement with our sequence analysis (**Fig. 5**), strong conservation was observed over most of the ClpC NTD surface (**Fig. 6B**), including around two putative phosphoarginine binding sites (46, 47). By contrast, surface conservation was lower and more localized on models of ClpB and ClpI NTDs (**Fig. 6A, C**). The high surface conservation on the ClpC NTD reinforces the idea that this module has multiple conserved interaction partners that contribute to the cellular function and regulation of ClpC enzymes.

**Figure 6.**
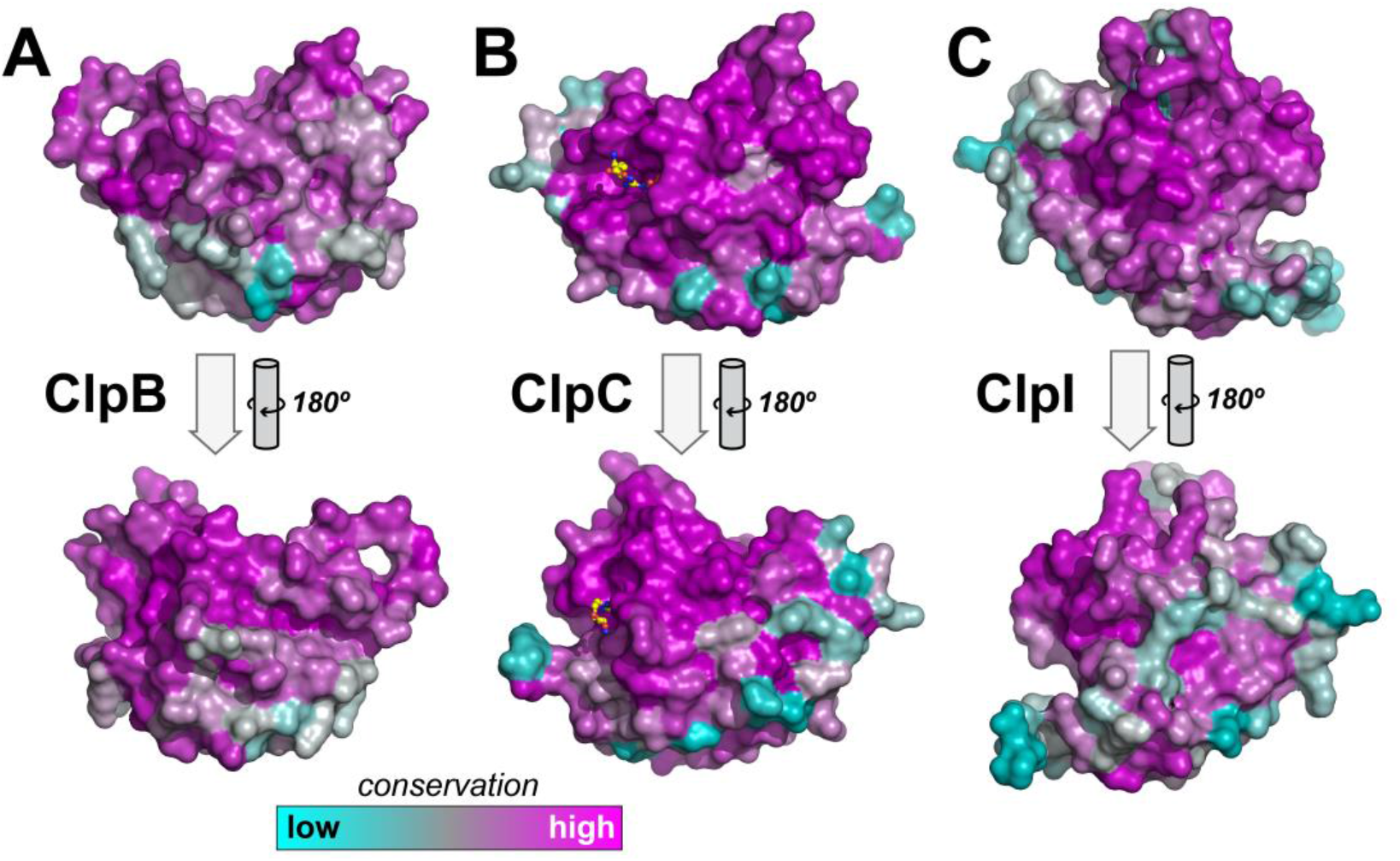
Surface conservation of Clp unfoldase NTDs. **A**) ClpB sequence conservation was projected onto the NTD surface from the AlphaFold model AF-P9WPD1-F1 of *M. tuberculosis* ClpB. **B**) ClpC sequence conservation was projected onto the surface of the crystal structure *M. tuberculosis* ClpC NTD (PDB ID: 3WDB). Phosphoarginine, shown as sticks, was modeled into conserved binding sties (46) as observed in the *B. subtilis* NTD, PDB ID: 5HBN. **C**) ClpI sequence conservation was projected onto the NTD from the AlphaFold model AF-A0A4P6UA74-F1 of *Rhodococcus sp*. ABRD24 ClpIa.

## DISCUSSION

Double-ringed Hsp100/Clp family unfoldases are ubiquitous in Actinomycetota. Based on sequence annotations and prior studies, Actinomycetota were known to possess two groups with distinct roles in protein homeostasis: ClpB enzymes that function as independent unfoldases and disaggregases (23), and ClpC enzymes that cooperate with the ClpP1P2 peptidase to carry out regulated proteolysis of folded proteins (4, 5, 7, 8, 12, 47, 48). An ability to confidently sort homologs into discrete enzyme families is critical for unraveling their individual contributions to cellular physiology. However, our initial sequence analysis of Actinomycetota ClpC homologs revealed numerous enzymes with similarity to ClpC and ClpB, but possessing sequence features that defied simple classification into these established groups. Our analysis here reveals the existence of a third evolutionarily distinct category of double-ringed Clp enzymes in Actinomycetota, which we term ClpI.

ClpI orthologs are less abundant than ClpB and ClpC enzymes, and appear in only a subset of Actinomycetota orders. Indeed, ClpI enzymes are absent in some of the more extensively studied species in Actinomycetota, including *Mycobacterium tuberculosis*, *Corynebacterium glutamicum*, and *Streptomyces coelicolor*, which may explain why they have not been previously described. Where they do occur, ClpI enzymes appear to exist alongside ClpB and ClpC. Accordingly, ClpI orthologs likely expand the complexity of protein quality control programs in these cells, providing additional points of regulation.

Interestingly, ClpI enzymes are divisible into subgroups, which likely have distinct cellular roles. The ClpIa subtype possess an LGF-loop similar in length to that of ClpC with an identifiable LGF-like motif, which is critical for docking of Clp unfoldases to the Clp peptidase (24, 40, 49, 50). ClpIa enzymes thus presumably serve as alternative unfoldase partners for ClpP1P2, and function in regulated proteolysis of client proteins. The region equivalent to the LGF-loop in the ClpIb subtype is short, ~5 amino acids, and lacks a conserved LGF-like motif. Thus, ClpIb enzymes likely operate as independent unfoldases, although we cannot rule out the possibility that they interact in an LGF-independent manner with ClpP2; with the ClpP1 ring, which has no known interaction partners to its outer face; or even with the 20S peptidase complex.

While ClpP-associated unfoldases have been shown to operate independently of ClpP in some contexts (51–55), to our knowledge, ClpI enzymes are the only group of Clp unfoldases that exist with and without LGF-motifs. This raises questions about whether these two subtypes possess differences in substrate preference or unfolding ability. An obligate unfoldase that works to rescue diverse misfolded proteins would likely benefit from more promiscuous substrate recognition than a ClpP-associated unfoldase, for which substrate selection must be tightly regulated to avoid harmful off-target proteolysis. Differences might also be expected in the degree of processively. Successful proteolysis of large and recalcitrant substrates is thought to require strong processively, permitting stepwise unfolding and translocation of over the course of many minutes (19, 50, 56–58). By contrast, the obligate unfoldase/disaggregase ClpB functions in a non-processive tug- and-release fashion under some circumstances, which may aid refolding of some targets (21, 59). *In vitro* biochemical studies of representative ClpI enzymes will help to answer these questions, and clarify their individual roles.

One other important finding, which arises from our improved ability to categorize ClpC orthologs, is the remarkable sequence conservation specifically among ClpC NTDs. The NTD is not known to possess enzymatic activity, nor to directly participate in substrate unfolding. Instead, it is thought to interact with substrates (eg, through phosphoarginine modifications (47, 60, 61)), with proteolytic adaptors (eg, the N-end rule adaptor ClpS (62)), and with the D1 ring and M-domain to influence the functional and oligomeric state of the entire unfoldase (61, 63, 64). Its conservation highlights the importance of these interactions across Actinomycetota. Moreover, it helps explain why multiple naturally occurring antibiotics have evolved to bind the ClpC NTD, and how antibiotics with overlapping binding sites can have different mechanistic effects on ClpC function (9, 46, 64, 65).

