## Supplemental Material for "Bioinformatic identification of ClpI, a class of Clp unfoldase in Actinomycetota"

**Table S1. Key residues that differ among ClpC/ClpB/ClpI enzymes.**

| ***B. subtilis* ClpC** | ***M. tuberculosis* ClpC1** | **Actinomycetota ClpC** | **Actinomycetota ClpB** | **Actinomycetota ClpI** |
|---|---|---|---|---|
| **S57** | L56 | L | P/Q | P |
| **T105** | T105 | T | T | P |
| **T160** | S168 | S | K | T |
| **D169** | N177 | N | D | D |
| **R191** | R199 | R | R | Q |
| **E194** | Q202 | Q | Q | E |
| **N298** | S306 | S | N | N |
| **Q310*** | Q318 | Q | R | H/R |
| **E397** | E405 | E | E | Q |
| **V431** | I439 | I | Q/E | V |
| **Q434** | Q442 | Q | G | E |
| **E446** | E454 | E | E | I |
| **M502** | M510 | M | M/L | L |
| **I542** | I550 | I | L | L |
| **L544** | A552 | A | L | L |
| **G599** | G607 | G | G | A |
| **Y611** | F619 | F | Y | Y |
| **E707*** | D715 | D | D/E | E |
| **H712*** | H720 | H/P | D/S/H | R/H |
| **E774*** | E782 | E | G | D |

* Noted as differing among proteobacterial ClpA/E/L/C proteins (66).

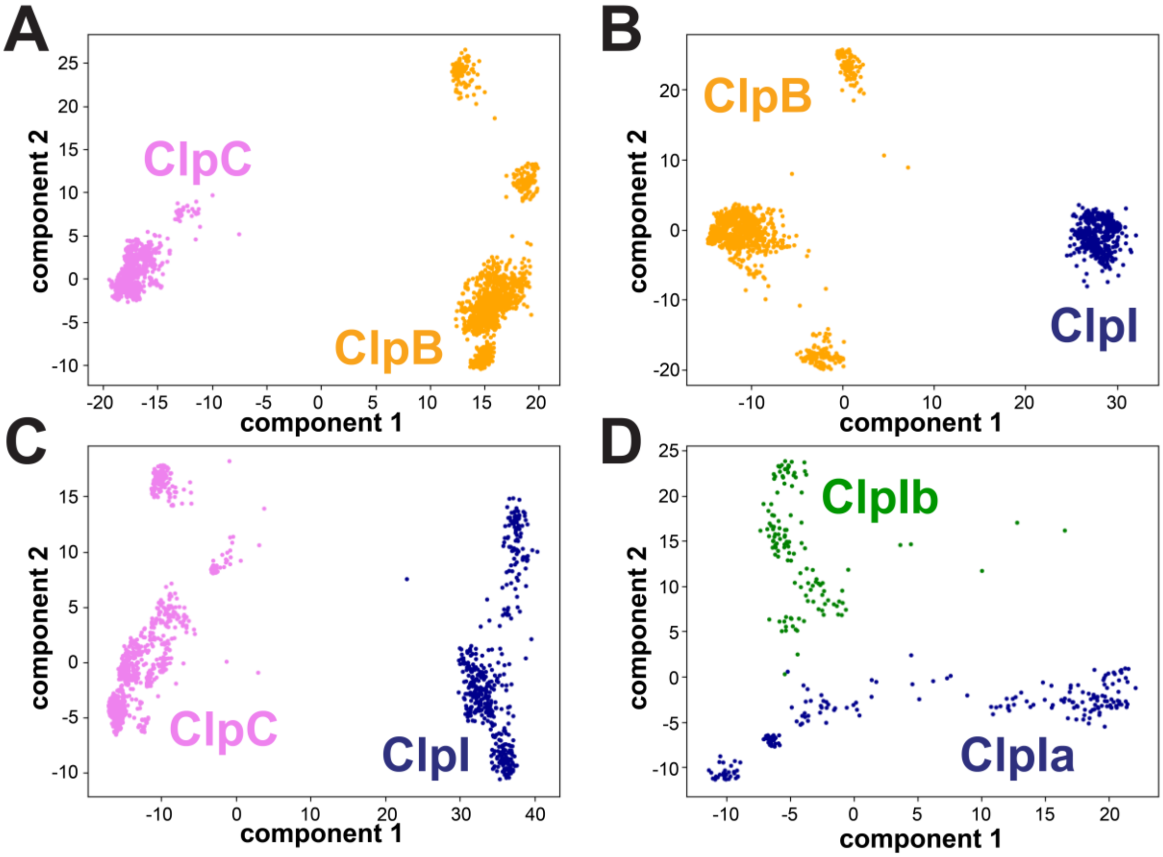


**Figure S1. Principal component analysis.** Clp enzyme orthologs from Actinomycetota were subjected to PCA analysis, comparing **A**) ClpC (violet) and ClpB (orange), **B)** ClpB (orange) and ClpI (blue), **C**) ClpC and ClpI, or **D)** ClpIa (blue) and ClpIb (green).

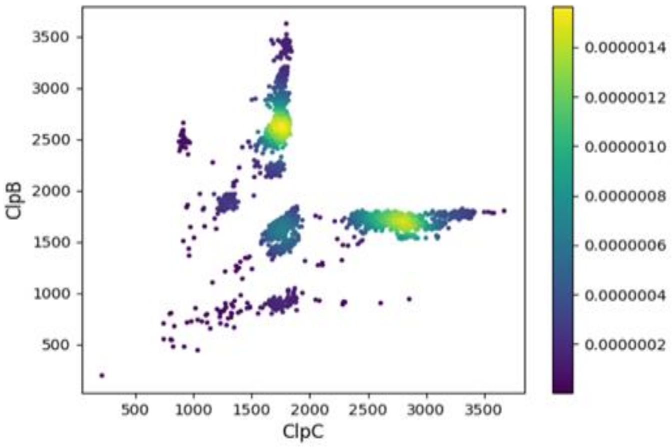


**Fig. S2. BLAST analysis of Actinomycetota ClpB/C orthologs omitting NTD.** The NTD region was removed from ClpC/B orthologs, and the resulting sequences were compared to *Mycobacterium smegmatis* ClpB (A0QQF0_MYCS2) and ClpC (CLPC1_MYCS2) references as in **Figure 2**. The *x* axis represents the BLAST score against the ClpC reference, and the *y* axis represents the BLAST score against the ClpB reference.
